# Cyclic di-GMP Increases Catalase Production and Hydrogen Peroxide Tolerance in *Vibrio cholerae*

**DOI:** 10.1101/631275

**Authors:** Nicolas L. Fernandez, Christopher M. Waters

## Abstract

*Vibrio cholerae* is a Gram-negative bacterial pathogen that causes the disease cholera, which affects nearly 1 million people each year. In between outbreaks, *V. cholerae* resides in fresh and salt water environments where it is able to persist through changes in temperature, oxygen, and salinity. One key characteristic that promotes environmental persistence of *V. cholerae* is the ability to form multicellular communities, called biofilms, that often adhere to biotic and abiotic sources. Biofilm formation in *V. cholerae* is positively regulated by the dinucleotide second messenger cyclic dimeric guanosine monophosphate (c-di-GMP). While most research on the c-di-GMP regulon has focused on biofilm formation or motility, we hypothesized the c-di-GMP signaling network encompassed a larger set of effector functions than reported. We found that high intracellular c-di-GMP increased catalase activity approximately 4-fold relative to strains with unaltered c-di-GMP. Genetic studies demonstrated that c-di-GMP mediated catalase activity was due to increased expression of the catalase encoding gene *katB*. Moreover, c-di-GMP mediated regulation of catalase activity and *katB* expression required the c-di-GMP dependent transcription factors VpsT and VpsR. Lastly, we found that high c-di-GMP increased survival after H_2_O_2_ challenge in a *katB, vpsR*, and *vpsT* dependent manner. Our results indicate antioxidant production is regulated by c-di-GMP in *V. cholerae* uncovering a new node in the growing VpsT and VpsR c-di-GMP signaling network.

**Importance:** As a result of infection with *V. cholerae*, patients become dehydrated leading to death if not properly treated. The marine environment is the natural reservoir for *V. cholerae* where it can survive alterations in temperature, salinity, and oxygen. The second messenger molecule c-di-GMP is an important signal regulating host and marine environmental persistence because it controls whether *V. cholerae* will form a biofilm or disperse through flagellar motility. In this work, we demonstrate another function of c-di-GMP in *V. cholerae* biology: promoting tolerance to the reactive oxygen species H_2_O_2_ through differential regulation of catalase expression. Our results suggest a mechanism where c-di-GMP simultaneously controls biofilm formation and antioxidant production, which could promote persistence in human and marine environments.

## Introduction

The Gram-negative bacterium *Vibrio cholerae* is the human pathogen that causes the diarrheal disease cholera. The most common route to infection is consumption of contaminated food or water, after which *V. cholerae* traverses the stomach and colonizes the small intestines. Cholera patients lose liters of fluid and dissolved ions through toxin-mediated changes to the host intestinal tract, allowing *V. cholerae* to exit the host, re-enter a water source, and perpetuate its infectious cycle. In addition to the harsh conditions of the human gastrointestinal tract, *V. cholerae* must adapt to numerous stresses in the marine environment. These environmental stresses include temperature fluctuations, eukaryotic predation, and exposure to chemical insults like reactive oxygen species (ROS) (1).

As an aquatic organism, *V. cholerae* is exposed to varying concentrations of dissolved oxygen and ROS produced abiotically through photochemical reactions between sunlight and dissolved organic matter in the ocean (2, 3). ROS can also be produced biotically through metabolic processes in aerobic environments by phytoplankton, another potential reservoir of *V. cholerae* (4). In response to the multiple routes of exposure to ROS, it is not surprising that *V. cholerae* has multiple ROS defense systems including two paralogues of the oxidative stress responsive transcription factor OxyR, two catalases, and multiple peroxidases (5–7). Another mechanism to increase tolerance to ROS is the production of surface adhered communities encased in an exopolysaccharide matrix also known as biofilms.

Many bacterial species, including *V. cholerae*, have increased tolerance to ROS such as hydrogen peroxide (H_2_O_2_) when grown in biofilms compared to planktonic counterparts (8–10). Biofilm formation in *V. cholerae* is regulated by the bacterial second messenger molecule cyclic dimeric guanosine monophosphate (c-di-GMP), which is produced by diguanylate cyclase (DGC) enzymes. C-di-GMP alters bacterial physiology by modulating transcription, translation, and/or protein function (11). In *V. cholerae*, a common mechanism of c-di-GMP signaling is modulation of gene expression by three c-di-GMP dependent transcription factors: VpsR, VpsT, and FlrA (12–14). c-di-GMP activates the transcription factors VpsR and VpsT, resulting in increased transcription of genes involved in synthesis of the biofilm matrix component *Vibrio* polysaccharide (VPS) (12, 13, 15). In contrast, c-di-GMP acts as an anti-activator of FlrA, which causes decreased expression of genes necessary for flagellar biosynthesis (14). C-di-GMP can also bind to two riboswitches, Vc1 and Vc2. Binding of c-di-GMP to Vc1 functions as an ON-switch to induce production of the adhesin GbpA, while binding of c-di-GMP to Vc2 functions as an OFF-switch to inhibit production of the transcription factor TfoY (16, 17). As cells receive signals to disperse from the biofilm, phosphodiesterase (PDE) enzymes, which degrade c-di-GMP, become activated. These PDEs then deplete the intracellular concentration of c-di-GMP, promoting a switch from a sessile biofilm lifestyle to a motile one (reviewed in (11)).

While VpsR and VpsT were initially discovered as regulators of biofilm production, other c-di-GMP dependent functions have emerged. For example, c-di-GMP and VpsR transcriptionally regulate genes in the type II secretion operon as well as *tfoY*, a gene involved in driving dispersive motility and regulating type VI secretion (17–19). VpsT negatively regulates expression of genes involved in flagellar biosynthesis; however, the mechanism is not known (13). Additionally, Ayala *et al*. demonstrated VpsT negatively regulates the transcription of the stationary phase sigma factor RpoS (20). Recently, we found that c-di-GMP and VpsT induced expression of the DNA repair gene *tag* to promote survival after alkylation stress (21). These studies demonstrate c-di-GMP regulation extends beyond biofilm formation and motility in *V. cholerae*.

In this study, we uncovered an additional role for c-di-GMP: positively regulating catalase activity by increasing transcription of the catalase encoding gene *katB* via a VpsR and VpsT dependent mechanism. We further show that c-di-GMP dependent catalase activity was necessary for survival after exposure to the ROS H_2_O_2_. Our results expand the regulatory network of c-di-GMP to include antioxidant production, demonstrating that elevated c-di-GMP enhances the oxidative stress response in *V. cholerae*.

## Materials and Methods

### DNA manipulations and growth conditions

*V. cholerae* C6706 Str2 was used as the wild-type and the low biofilm forming derivative Δ*vpsL* was used as the Parent strain in the text (22, 23). All vectors were constructed by Gibson Assembly (NEB). Chromosomal deletion strains were constructed using the allele exchange vector pKAS32 digested with KpnI and SacI (NEB, High Fidelity). Luciferase reporters were constructed using the luciferase reporter vector pBBRlux digested with BamHI and SpeI (NEB). Expression vectors for VpsT and VpsR were constructed by removing the ribosome binding site (RBS), green fluorescent protein, and chloramphenicol acetyltransferase from pEVS143 with BamHI and EcoRI digests (NEB) (23). The VpsT purification vector was constructed elsewhere (21). Expression vectors for *katB* and *katG* were constructed as follows: pHERD20T was amplified with primers flanking the ampicillin resistance gene using inverse PCR resulting in a linear PCR fragment lacking the ampicillin resistance gene (24). The chloramphenicol resistance gene from pBBRlux was amplified by PCR and the two linear fragments were circularized by Gibson Assembly. Plasmids were introduced into *V. cholerae* through biparental conjugation using *Escherichia coli* S17 as the donor strain. *V. cholerae* harboring the plasmid of interest was selected for using Polymixin B (10 U/mL) with the relevant antibiotic. Antibiotics and reagents were used at the following concentrations unless otherwise stated: Ampicillin (100 μg/mL), Kanamycin (100 μg/mL), chloramphenicol (10 μg/mL), and 100 μM isopropyl β-D-1-thiogalactopyranoside (IPTG). Cultures were grown in Lysogeny Broth (LB, Acumedia) at 35°C, 220 RPM unless otherwise stated.

### Catalase assay

Measurement of catalase activity was adapted from (25) with the following changes: Overnight cultures were diluted to a starting OD_600_ of 0.04 in 5 mL LB in 18 × 150 mm borosilicate test tubes supplemented with necessary antibiotics and IPTG. Cultures were grown at 35°C with shaking at 220 RPM until the OD_600_ reached approximately 2.0, moved to 15 mL falcon tubes (Corning®), and were pelleted by centrifugation (4,000 × g for 3 minutes). Pellets were resuspended in 100 μL of sterile 1X PBS to create a viscous cell solution that were adjusted to an OD_600_ of 150 in 100 μL final volume in Pyrex test tubes (13 × 100 mm, borosilicate). 200 μL of catalase reaction buffer (1% Triton-X100, 15% hydrogen peroxide in 1X PBS) was added to the test tubes and the solution was mixed using disposable 10 μL loops (BD Difco). Tubes were incubated at room temperature until gas production subsided (approximately 5-10 minutes). A standard curve was generated by mixing purified bovine catalase (Sigma, 570 U/μL) diluted in 1X PBS with 200 μL of catalase reaction buffer. At 10 minutes of incubation, images of the tubes were taken with an iPad Air (iOS 12.1.4) and the height from the bottom of the tube to the top of the foam were measured in both the standards and samples using the software ImageJ and an internal 1-inch reference mark for each picture. GraphPad Prism was used to generate the standard curve and interpolate the sample catalase activity using linear regression. Data presented is catalase activity (Units) normalized to cell number (OD_600_).

### RNA Isolation and quantitative real-time PCR (qPCR)

Three biological replicate overnight cultures were diluted to a starting OD_600_ of .04 in 2 mL LB supplemented with ampicillin and IPTG and grown until an OD_600_ of approximately 1.0 at 35°C and 220 RPM. 1 mL of each replicate was pelleted and RNA was extracted using the TRIzol® reagent following the directions in the manual (Thermo Fischer Scientific). Purified DNA was quantified using a NanoDrop spectrophotometer (Thermo Fischer Scientific). 5 μg of purified RNA was treated with DNAse (Turbo DNAse, Thermo Fischer Scientific). cDNA synthesis was carried out using the GoScript™ Reverse Transcription kit (Promega). cDNA was diluted 1:30 into molecular biology grade water and used as template in qRT-PCR reactions using SYBR Green (Applied Biosystems™) as the method of detection. Reactions consisted of 5 μL 2.5 μM primer 1, 5 μL of 2.5 μM primer 2, 5 μL of diluted cDNA template, and 15 μL of 2X SYBR green (see Table S4 for primer sequences). Each plate had technical duplicates and biological triplicate samples as well as no reverse transcriptase controls to check for genomic DNA contamination. The StepOnePlus Real Time PCR system was used for qRT-PCR with the following thermocycling conditions: 95°C for 20 seconds then 40 cycles of 95°C for 2 seconds and 60°C for 30 seconds. Melting curves were included to ensure PCR products had single amplicons and primer dimers were absent. Data was analyzed by the ΔΔCt method using *gyrA* as a housekeeping or reference target.

### Luciferase reporter assays

#### A) *V. cholerae* reporter assays

Overnight cultures of *V. cholerae* harboring *katB* transcriptional fusions to luciferase in pBBRlux were diluted 1:100 in 1 mL LB supplemented with ampicillin, chloramphenicol, and IPTG. 200 μL of cell solution was aliquoted into wells of a black 96-well plates (Costar). Plates were incubated at 35°C with shaking at 220 RPM until the OD_600_ reached approximately 0.25, and luciferase activity was measured using an Envision plate reader (Perkin Elmer). Luciferase activity (RLU) was normalized for cell number by dividing RLU by the OD_600_ at the time of the reading (Normalized Luminescence). For experiments where H_2_O_2_ were added to the cultures, overnight cultures of *V. cholerae* were diluted as described above except H_2_O_2_ was added to cultures at a final concentration of 50 μM when OD_600_ values reached approximately 0.225, followed by shaking at 35°C for an additional 30 minutes before measuring luciferase and OD_600_.

#### B) *E. coli* DH10b luciferase assays

Overnight cultures of *E. coli* DH10b harboring vectors to modulate transcription factor, c-di-GMP production, and the luciferase reporter were diluted 1:100 as described above and grown at 35°C, 220 RPM until the OD_600_ reached 0.45. Luciferase activity was measured and normalized to the OD_600_ to yield normalized luminescence.

### Protein purification and electrophoretic mobility shift assays (EMSA)

C-terminal HIS-tagged VpsT purification and EMSAs experiments using the FAM labeled *katB* promoter region from *katB2* were carried out as previously described (21). For purification, an overnight culture of *E. coli* BL21 harboring the pET28b-VpsT expression construct was diluted 1:100 into 250 mL LB supplemented with kanamycin in a 1 L flask. The culture was grown until an OD_600_ of approximately 0.7 at which point 1 mM IPTG was added and the culture conditions were shifted to 16°C with shaking at 160 RPM for 16 hours to induce protein production. Protein purification was carried out by standard Ni-NTA resin purification protocols (19). For EMSA experiments, varying concentrations of purified HIS-tagged VpsT (0 – 600 nM) were incubated with 2.5 nM FAM-labeled *katB* probe in VpsT buffer (25 mM Tris-Cl, 150 mM NaCl, 5 mM β-mercapthol, pH – 7.5) at 30°C for 30 minutes. The binding reaction was loaded into pre-run 5% non-denaturing TBE gels and gel electrophoresis was done by applying 90 volts for 90 minutes at 4°C. Fluorescent detection and images of the gels and were taken using a Typhoon FLA 9000 imager and the requisite software (GE Healthcare Life Sciences).

### Hydrogen peroxide survival assay

Overnight cultures were diluted to a starting OD_600_ of 0.04 in 1 mL LB supplemented with necessary antibiotics and IPTG in 1.5 mL microcentrifuge tubes. 140 μL aliquots were added to a 96-well plate (Costar) and grown until an OD_600_ of 0.3. H_2_O_2_ solutions were made from fresh H_2_O_2_ stocks in light-impermeable microcentrifuge tubes and sterile 1X PBS. At time 0, 10 μL of H_2_O_2_ was added to the cell solution and growth was monitored over time by measuring OD_600_.

### Measurement of intracellular c-di-GMP

Overnight cultures of Δ*vpsL* harboring pBRP1 (QrgBMut) were diluted 1:100 in 2 mL LB ampicillin in 18 × 150 mm borosilicate test tubes and were grown to an OD_600_ of 1.0. The cultures were split into two 1 mL aliquots in microcentrifuge tubes, and H_2_O_2_ was added to one aliquot at a final concentration of 500 μM. An equal volume of water was added to the other aliquot as the untreated control. Cultures were incubated statically at room temperature for 30 minutes and collected for total protein quantification and nucleotide extraction. Briefly, 100 μL of culture was removed from each tube to quantify total protein, pelleted by centrifugation at full speed (15,000 × g) for 1 minute, resuspended in 100 μL 1X PBS with 10% sodium dodecyl sulfate (SDS), and boiled at 95°C for 10 minutes. Lysed cell solutions were centrifuged at 15,000 × g for 1 minute and the supernatant was removed and placed in new tubes. Total protein was quantified using the DC Protein Assay (Bio-Rad) following the instructions in the manual. Protein standards consisting of bovine serum albumin (provided in DC Protein Assay) were used to generate a standard curve to interpolate unknown sample concentrations. Nucleotide extractions were carried out following the protocol here (26) with the following changes. 900 μL of the remaining culture were pelleted at 15,000 × g for 1 minute in a benchtop microcentrifuge. The supernatants were removed, and the remaining pellets were resuspended in 100 μL nucleotide extraction buffer (40:40:20 methanol/acetonitrile/water with 0.1 N formic acid). The extraction solution was incubated at 20°C for 20 minutes and pelleted for 10 minutes at 15,000 × g. The supernatants were placed into new microcentrifuge tubes, and the solutions were dried overnight using a heated, vacuum centrifuge (SpeedVac Concentrator, Savant). The resulting dried pellets were resuspended in 100 μL HPLC-grade water and subjected to mass spectrometry analysis for quantification of c-di-GMP (27). Data is represented as pmol of c-di-GMP normalized by total cellular protein (mg).

### Statistical Analysis

Data are represented as the mean ± SD. Statistical analyses (details in figure legends) were calculated with GraphPad Prism Ver. 6 (GraphPad, San Diego, CA). A p-value of < 0.05 was considered statistically significant.

## Results

### C-di-GMP Positively Regulates Catalase Activity

We have shown that c-di-GMP regulates genes involved in DNA repair and that this regulation increased tolerance to the methylating agent methyl methanesulfonate (MMS) (21). Thus, we hypothesized c-di-GMP had a role in mitigating other forms of cellular stress besides DNA methylation damage. We chose to test if c-di-GMP increased H_2_O_2_ tolerance because it is a common ROS produced by aerobic microorganisms as a byproduct of cellular respiration, and H_2_O_2_ is found in high concentrations in marine environments (reviewed in (28)). To test the effects of c-di-GMP on H_2_O_2_ tolerance in *V. cholerae*, we expressed the *Vibrio harveyi* DGC QrgB in a strain unable to form mature biofilms (Δ*vpsL*, designated as the parent strain for this text). We used this strain because expressing QrgB in WT *V. cholerae* induces biofilm production, which would introduce a confounding variable in our experiments (23). As a control, we expressed an inactive allele of QrgB (QrgB^Mut^) which is unable to synthesize c-di-GMP. Immediately after the addition of H_2_O_2_, both cultures began to produce gas bubbles, which is a phenomenon indicative of catalase activity as H_2_O_2_ is degraded into water and gaseous oxygen. Interestingly, the culture with higher intracellular c-di-GMP exhibited a higher amount of gas production despite the cultures having similar numbers of bacteria, suggesting c-di-GMP increased catalase activity. Quantification of catalase activity revealed an approximate 5-fold increase when comparing strains expressing QrgB to QrgB^Mut^ (Figure 1), indicating c-di-GMP increased catalase activity. Importantly, we have previously demonstrated that the concentration of c-di-GMP generated by QrgB overexpression is similar to that seen naturally in the low-cell-density quorum sensing state, demonstrating that these results are physiologically relevant (17).

**Figure 1.**
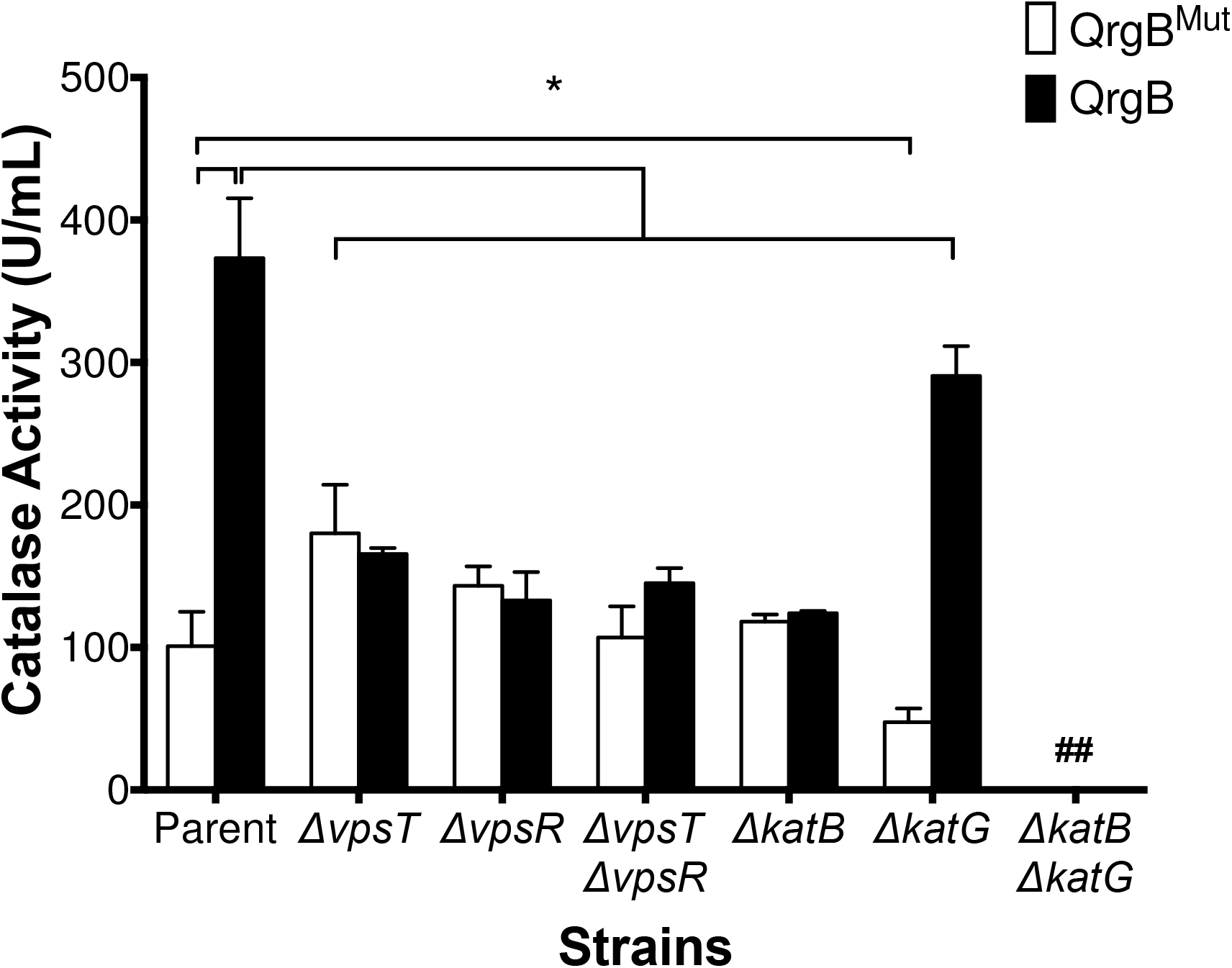
C-di-GMP increases catalase activity. Catalase activity was measured in the parent and mutant strains overproducing QrgB^Mut^ (white bars) or QrgB (black bars). Brackets and * indicate differences with a p < .05 determined by Two-Way ANOVA followed by Tukey’s multiple comparison test. Data are the average of three biological replicates with error bars indicating standard deviation. ## indicates activity was below the limit of detection.

c-di-GMP can directly modulate protein activity or change gene expression through allosteric interactions with c-di-GMP-dependent transcription factors or riboswitches (13–15). The c-di-GMP dependent transcription factors VpsT and VpsR induce transcription of genes involved in biofilm formation, protein secretion, and DNA repair in high c-di-GMP conditions (12, 13, 17, 18, 21). Therefore, we hypothesized that these transcription factors control c-di-GMP regulated catalase activity. To test this hypothesis, we repeated the catalase assay using Δ*vpsT*, Δ*vpsR*, and the Δ*vpsT*Δ*vpsR V. cholerae* mutants and observed a loss of c-di-GMP mediated induction of catalase activity, suggesting the increased catalase activity was part of the VpsR/VpsT/c-di-GMP regulatory network (Figure 1).

*V. cholerae* encodes two catalases: *katG* (VC1560), a bifunctional enzyme with both catalase and peroxidase functions, and *katB* (VC1585), which only exhibits catalase activity (7, 29). Mutants of these enzymes render *V. cholerae* more susceptible to H_2_O_2_ treatment; however, regulation of either *katB* or *katG* by c-di-GMP has not been described (29). We therefore measured c-di-GMP induction of catalase activity in the mutants Δ*katB*, Δ*katG*, or Δ*katB*Δ*katG*. We observed that the Δ*katB* background did not display c-di-GMP regulated catalase activity but did possess basal level catalase activity similar to the parent strain expressing QrgB^Mut^ (Figure 1). Expression of *katB* from an arabinose-inducible, multicopy plasmid in the Δ*katB* background complemented catalase activity regardless of the levels of c-di-GMP (Figure S1). Strains lacking *katG* were still able to induce catalase activity by c-di-GMP; however, the level of catalase activity induced by c-di-GMP was approximately 20% lower than that of the parent strain (Figure 1). Additionally, catalase activity in the Δ*katG* strain expressing QrgB^Mut^ was approximately 2-fold lower than the parent strain in the same condition (Figure 1). Strains lacking both *katG* and *katB* did not have measurable catalase activity under these conditions, which is expected because these are the only annotated genes encoding catalase activity in the El Tor *V. cholerae* reference genome N16961 (7) (Figure 1).

### Characterization of the *katB* Promoter and regulation of *katB* expression

Our results suggest *katB*, but not *katG*, is positively regulated by c-di-GMP at the transcriptional level. We addressed this hypothesis by measuring *katB* mRNA from cultures inducing QrgB and QrgB^Mut^ using qRT-PCR. We found that *katB* expression increased approximately 12-fold in strains expressing QrgB relative to QrgB^Mut^ (Figure 2).

**Figure 2.**
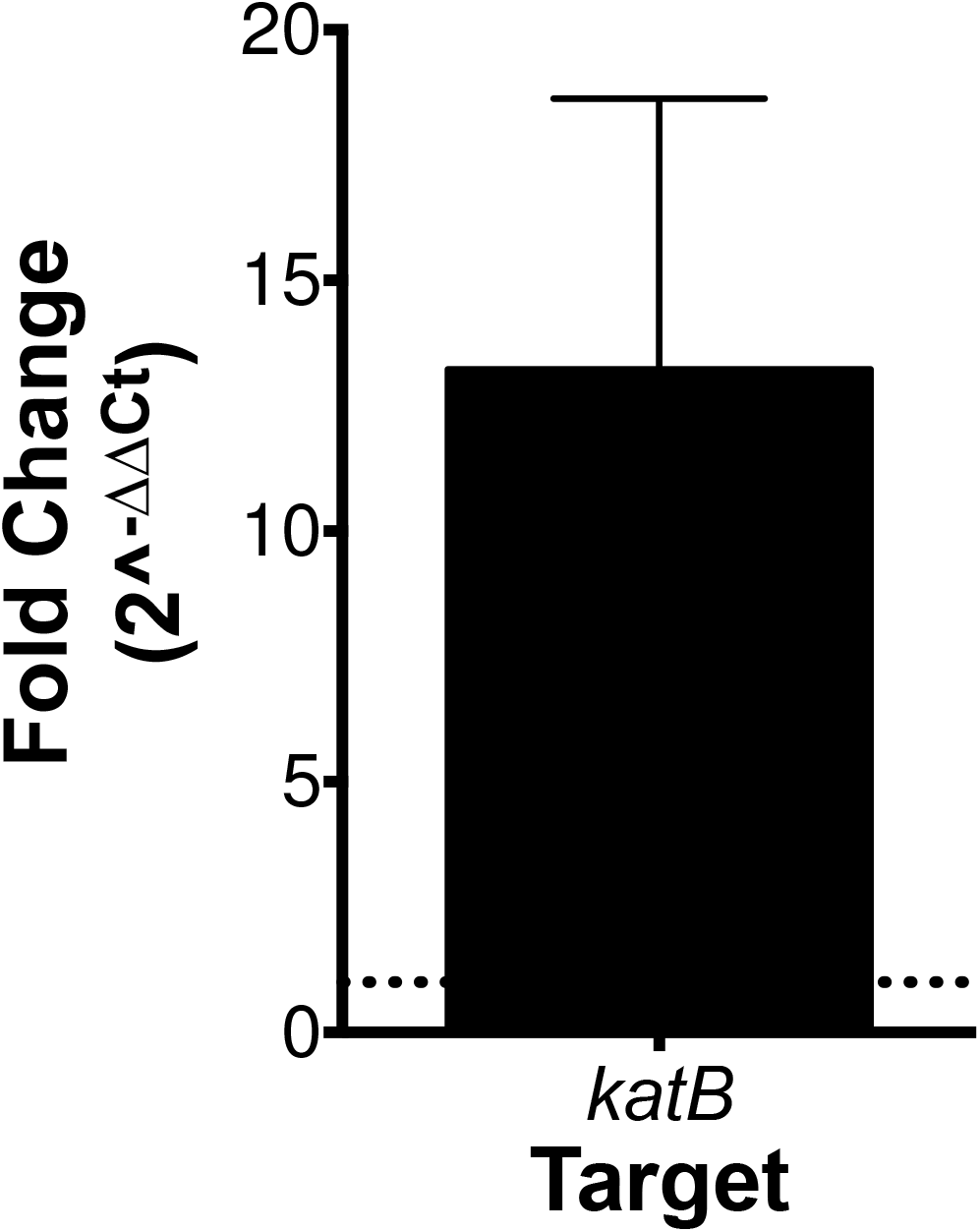
C-di-GMP increases *katB* mRNA abundance. qRT-PCR analysis comparing *katB* mRNA abundance between QrgB and QrgB^Mut^ conditions using the ΔΔCt method. Dashed horizontal line represents a fold change of 1. Bars represent the average of three biological replicates with error bars indicating standard deviation.

We next were interested in the promoter architecture of the region upstream of *katB* and hypothesized specific regions in the *katB* promoter were necessary for c-di-GMP mediated transcriptional control. We tested this hypothesis by measuring luciferase activity from a series of *katB* transcriptional reporters that have been truncated at the 5’ end in the presence of QrgB or QrgB^Mut^ expression (Figure 3A). In the full-length promoter construct *katB1*, QrgB induced *katB* expression 5-fold compared to strains over-producing QrgB^Mut^ (Figure 3A). Inclusion of 212 bp upstream of *katB* was sufficient to maintain c-di-GMP induction (Figure 3A, *katB2* and *katB3*). However, if the promoter was truncated to include 172 bp upstream of *katB (katB4*), c-di-GMP dependent induction of *katB* was abrogated. This result suggested the necessary cis-acting sequences for c-di-GMP mediated activation of *katB* expression are found between −172 and −212 bp upstream of the *katB* relative to the ATG start codon (Figure 3A). Deleting 30 bps from *katB4* resulted in expression levels similar to that of the promoter-less vector control regardless of c-di-GMP concentrations (*katB5*), suggesting components necessary for the basal expression of *katB* are between −172 and −142 relative to the ATG start codon (Figure 3A). We also measured the effect of c-di-GMP on a katG-luciferase transcriptional fusion and did not observe significant c-di-GMP dependent changes in expression, in agreement with our initial hypothesis (Figure 3B).

**Figure 3.**
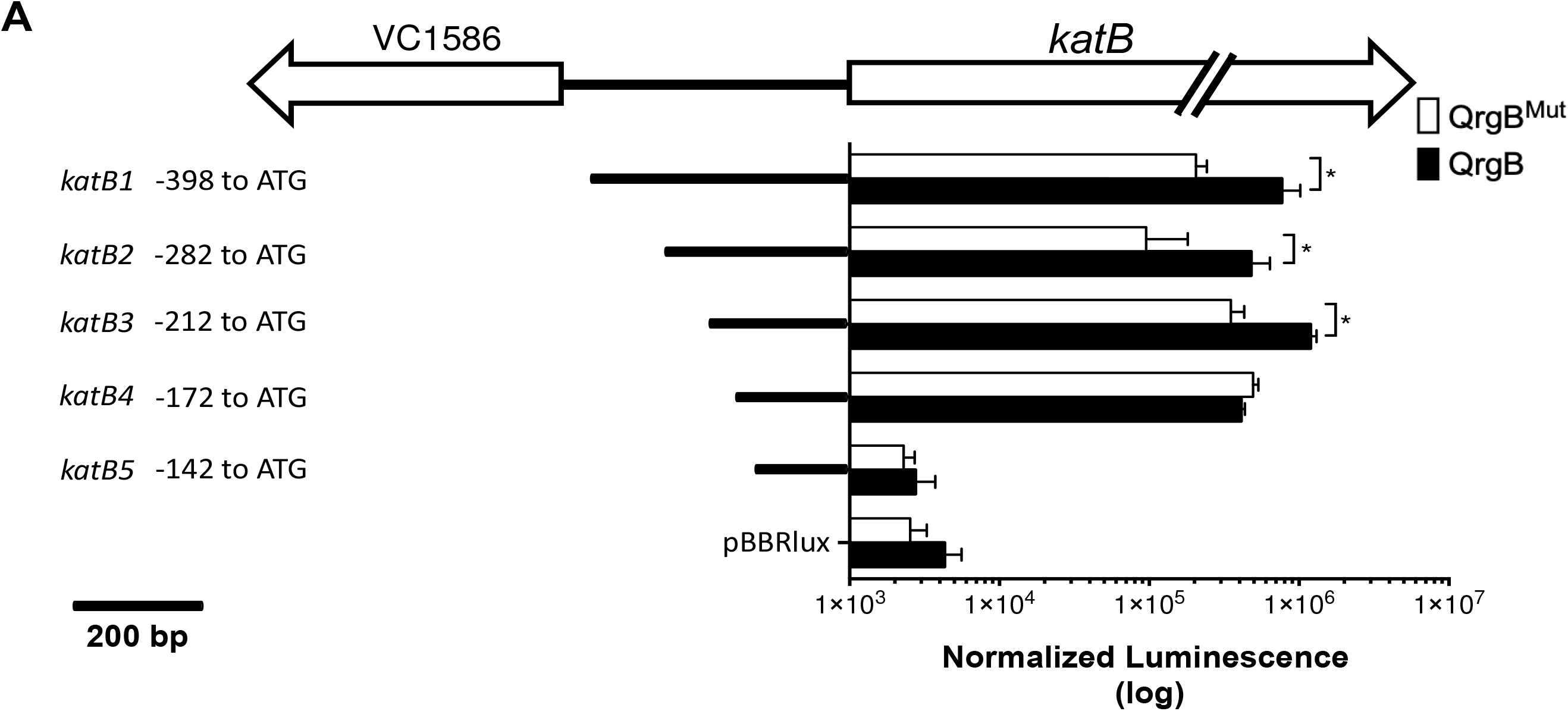
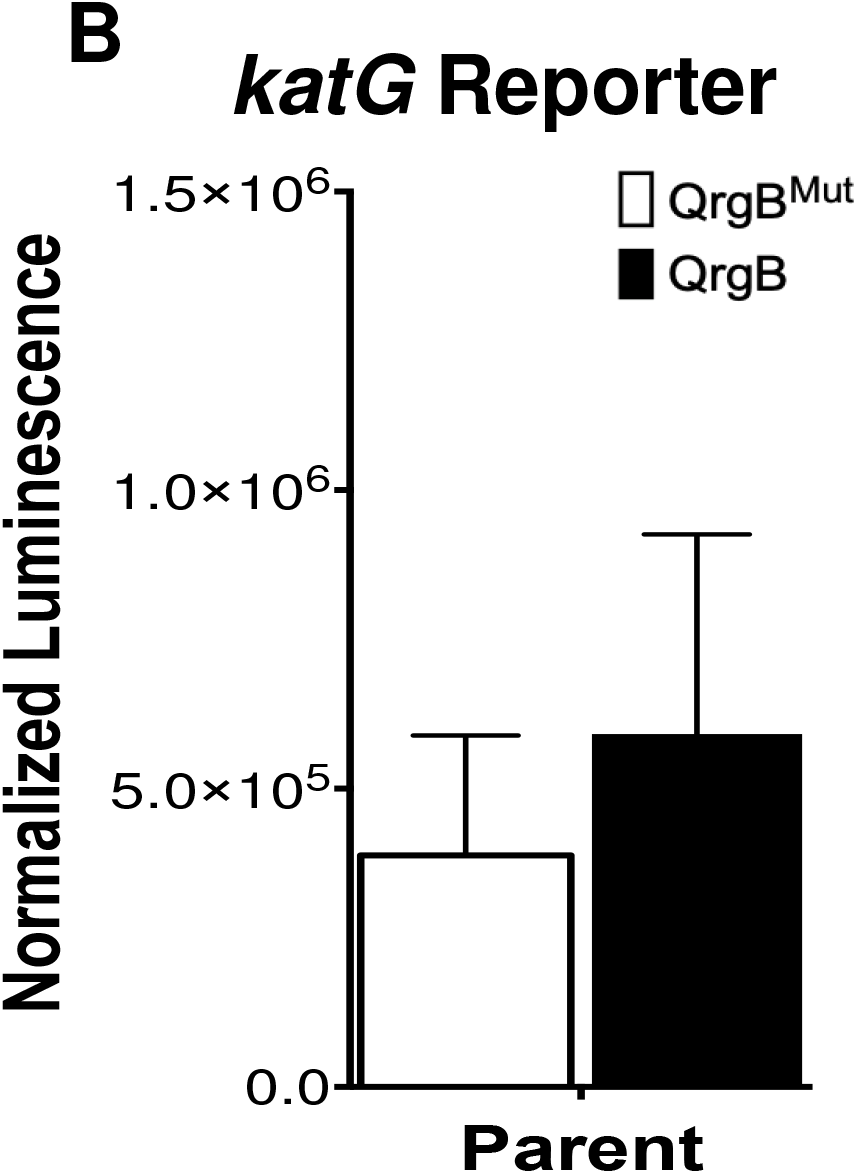
Characterization of the *katB* promoter. **A) *katB* genetic locus and promoter truncations and reporter activity.** 5’ promoter truncations were constructed and cloned upstream of the luciferase operon. Lines and numbers on the Y-axis indicate length of promoter relative to the ATG start codon for each promoter construct. Normalized luminescence is light production normalized to OD_600_ to control for cell number. For each promoter construct, luminescence was measured while overproducing QrgB^Mut^ (white bars) or QrgB (black bars). An empty pBBRlux vector control was included in each experiment. Brackets and * indicate comparisons with a p-value < .05 determined by Two-Way ANOVA followed by Tukey’s multiple comparisons testing. Data are the averages of three biological replicates and error bars indicate standard deviation. **B)** *katG* expression was measured in the parent strain while overproducing QrgB^Mut^ (white bars) or QrgB (black bars). Bars represent averages of three biological replicates with error bars indicating standard deviation.

VpsT and VpsR are necessary for the c-di-GMP-dependent induction of catalase activity, suggesting that these transcription factors activate transcription of *katB* at high intracellular concentrations of c-di-GMP (Figure 1). Thus, we hypothesized that c-di-GMP mediated induction of *katB* would be lost in strains lacking *vpsT, vpsR*, or both *vpsT* and *vpsR*. We tested this hypothesis by measuring *katB2* reporter activity under different c-di-GMP conditions in the parent, single, and double knockout strains. We found that *katB2* expression increased 5-fold in the parent background. *katB* expression increased 2.5 to 3-fold in the Δ*vpsT* and Δ*vpsR* backgrounds, but the differences in expression between QrgB^mut^ and QrgB were not statistically significant (Figure 4A). It was only in the double mutant Δ*vpsT*Δ*vpsR* background that c-di-GMP mediated *katB* expression was completely lost (Figure 4A). These data suggest that both VpsR and VpsT are needed for full induction by c-di-GMP.

**Figure 4.**
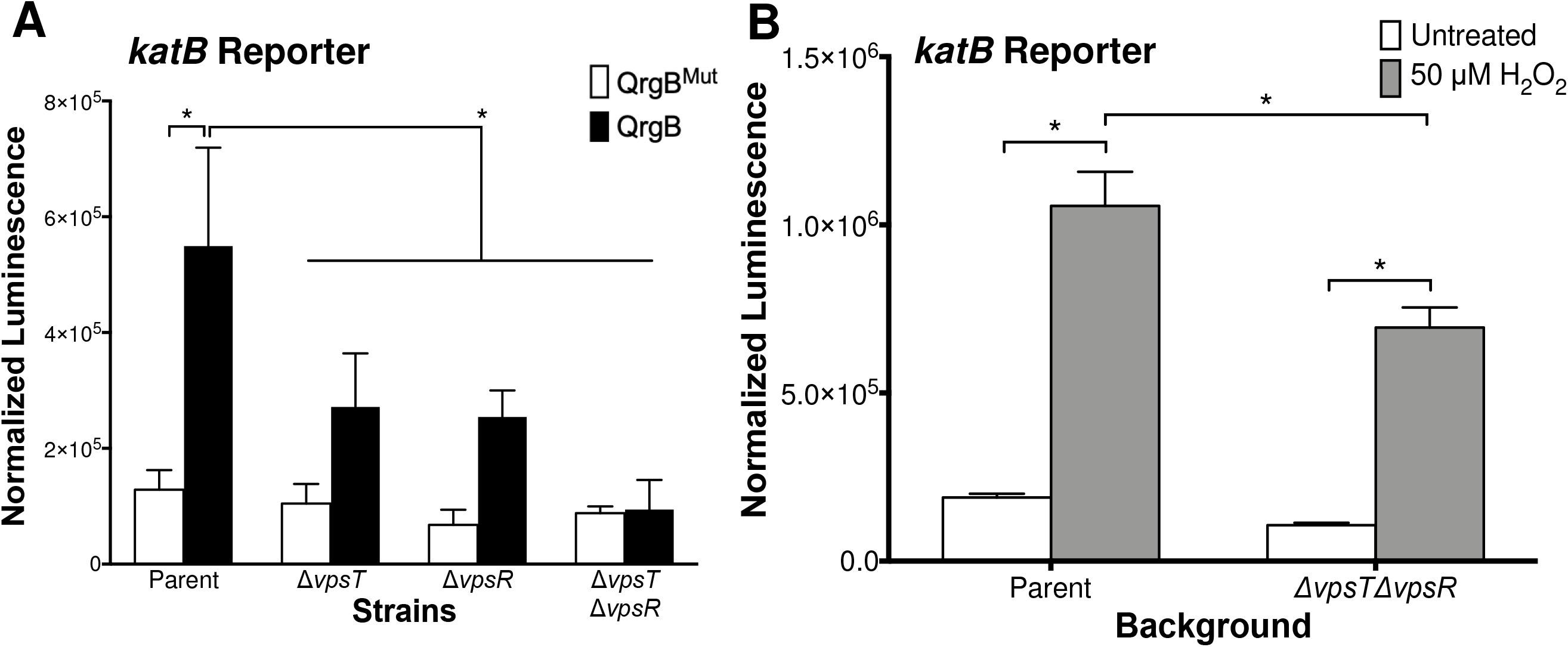
*vpsT* and *vpsR* are necessary for c-di-GMP mediated expression of *katB* but dispensable for H_2_O_2_ induction. **A)** *katB* expression was measured in parent, Δ*vpsT*, Δ*vpsR*, and Δ*vpsT*Δ*vpsR* backgrounds while overproducing QrgB^Mut^ (white bars) or QrgB (black bars). Brackets and * indicate differences with a p-value < .05 determined by Two-Way ANOVA followed by Tukey’s multiple comparison test. Data are average of three biological replicates. Error bars indicate standard deviation. **B)** *katB4* expression was measured in the parent and Δ*vpsT*Δ*vpsR* background 30 minutes after addition of 50 μM H_2_O_2_ or PBS control (untreated). Data are the average of three biological replicates and error bars indicate standard deviation.

*katB* expression is up-regulated in response to hydrogen peroxide in *V. cholerae* through the transcriptional activator OxyR (6, 29). To determine if the same region necessary for c-di-GMP mediated regulation of *katB* was necessary for H_2_O_2_ induction of *katB*, we measured the ability of H_2_O_2_ to induce *katB4*, the promoter region that no longer responded to c-di-GMP (Figure 3B). We found that 50 μM H_2_O_2_ induced *katB4* expression approximately 6-fold in the parent background (Figure 4B). Next, to determine if VpsT and VpsR contributed to the H_2_O_2_ inducible response of the *katB4* promoter, we repeated the assay in a Δ*vpsT*Δ*vpsR* background. In this background, H_2_O_2_ increased *katB4* expression to the same extent as the parent, however the basal and induced expression level was 1.5-fold lower compared to the parent (Figure 4B).

In *Mycobacterium smegmatis*, H_2_O_2_ can act as a first messenger to promote c-di-GMP synthesis (30). Whether H_2_O_2_ acts as a first messenger to modulate c-di-GMP in *V. cholerae* has not been demonstrated. To test the hypothesis that H_2_O_2_ acts as a first messenger, we measured intracellular c-di-GMP in control and H_2_O_2_ treated cultures and found no differences in intracellular c-di-GMP (Figure S2). Together, these results indicate that *katB* transcription is induced by c-di-GMP and the oxidative stress response through distinct regulatory mechanisms and demonstrate that H_2_O_2_ does not alter global levels of intracellular c-di-GMP.

### VpsT Induces *katB* Expression in a Heterologous Host and Binds to the *katB* Promoter In Vitro

To test whether VpsT or VpsR directly regulates *katB*, we used *Escherichia coli* as a heterologous host to measure *katB* expression at high versus low concentrations of c-di-GMP in the presence of either VpsR or VpsT. We reasoned the genetic dissimilarity between *V. cholerae* and *E. coli* would allow us to isolate the effects of VpsR and VpsT directly on the *katB* promoter without the effect of these transcription factors regulating each other’s expression (23, 31, 32). Expression of QrgB with an empty vector increased *katB* expression in *E. coli* approximately 2.5-fold when compared to expression of QrgB^Mut^ through an unknown mechanism (Figure 5). Similarly, co-expression of VpsR along with QrgB resulted in approximately the same fold change (2-fold) as the empty vector, indicating VpsR is not sufficient to induce *katB* expression when expressed alone in *E. coli* along with increased c-di-GMP (Figure 5). Expression of VpsT and QrgB, however, resulted in transcription from the *katB* promoter increasing 25-fold (Figure 5), suggesting that VpsT directly regulates *katB* transcription in response to c-di-GMP.

**Figure 5.**
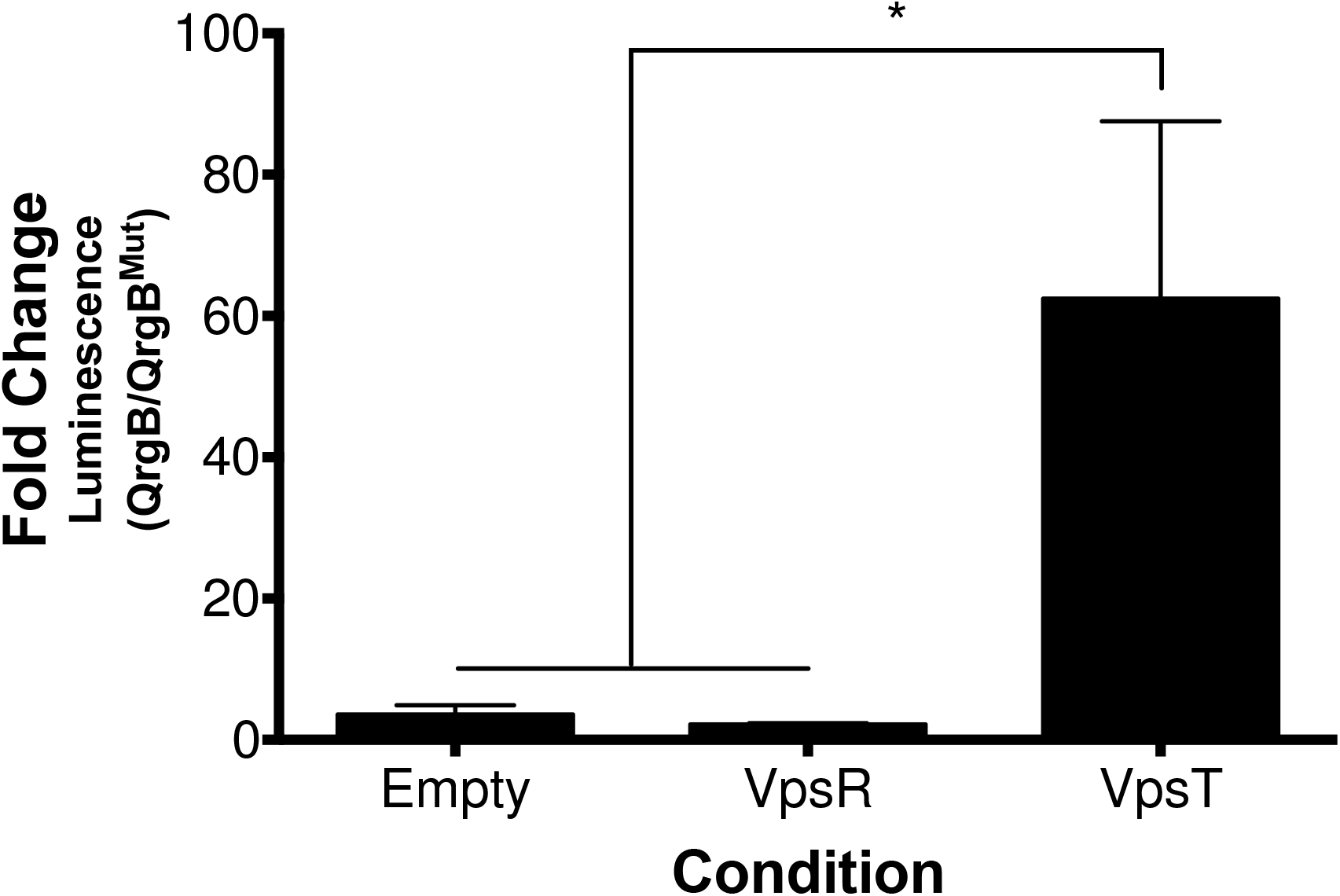
VpsT and c-di-GMP induce *katB* expression in a heterologous host. QrgB^Mut^ or QrgB were overproduced along with either an empty vector, VpsR, or VpsT in DH10b *E. coli* harboring the *katB3* promoter fused upstream of the luciferase operon. Data is the average fold change in luminescence between strains overproducing QrgB and QrgB^Mut^ from three biological replicates and error bars are standard deviation. Brackets and * indicate a statistically significant difference determined by One-Way ANOVA followed by Tukey’s multiple comparison testing.

As a transcription factor, VpsT binds to DNA in the presence of c-di-GMP to modulate gene expression (13, 20, 21, 32). Since VpsT was necessary for c-di-GMP mediated *katB* expression and was able to induce *katB* expression when expressed in a heterologous host, we hypothesized VpsT directly interacted with its promoter in a c-di-GMP-dependent manner. Thus, we purified C-terminal HIS-tagged VpsT and measured its ability to bind to the *katB2* promoter in vitro. VpsT only partially shifted the *katB2* probe in the absence of c-di-GMP at the highest concentration tested (600 nM) (Figure 6, lane 5). However, with the addition of 50 μM c-di-GMP, VpsT was able to decrease the intensity of the unshifted band at 150 nM and completely shift the probe at 300 nM (Figure 6, lanes 10-12). Addition of a 100-fold molar excess unlabeled 20bp oligonucleotide, composed of the VpsT binding site found in the *vpsL* promoter, was able to outcompete VpsT binding to the labeled *katB2* probe in the presence or absence of c-di-GMP (Figure 6, lanes 7, 14) (32). When the unlabeled competitor had transversion mutations introduced into the palindromic region, it was no longer able to abrogate the VpsT-*katB2* band migration (Figure 6, lanes 6, 13) (32). Together, the in vivo and in vitro data suggest VpsT directly interacts with the *katB* promoter to stimulate expression under high c-di-GMP conditions.

**Figure 6.**
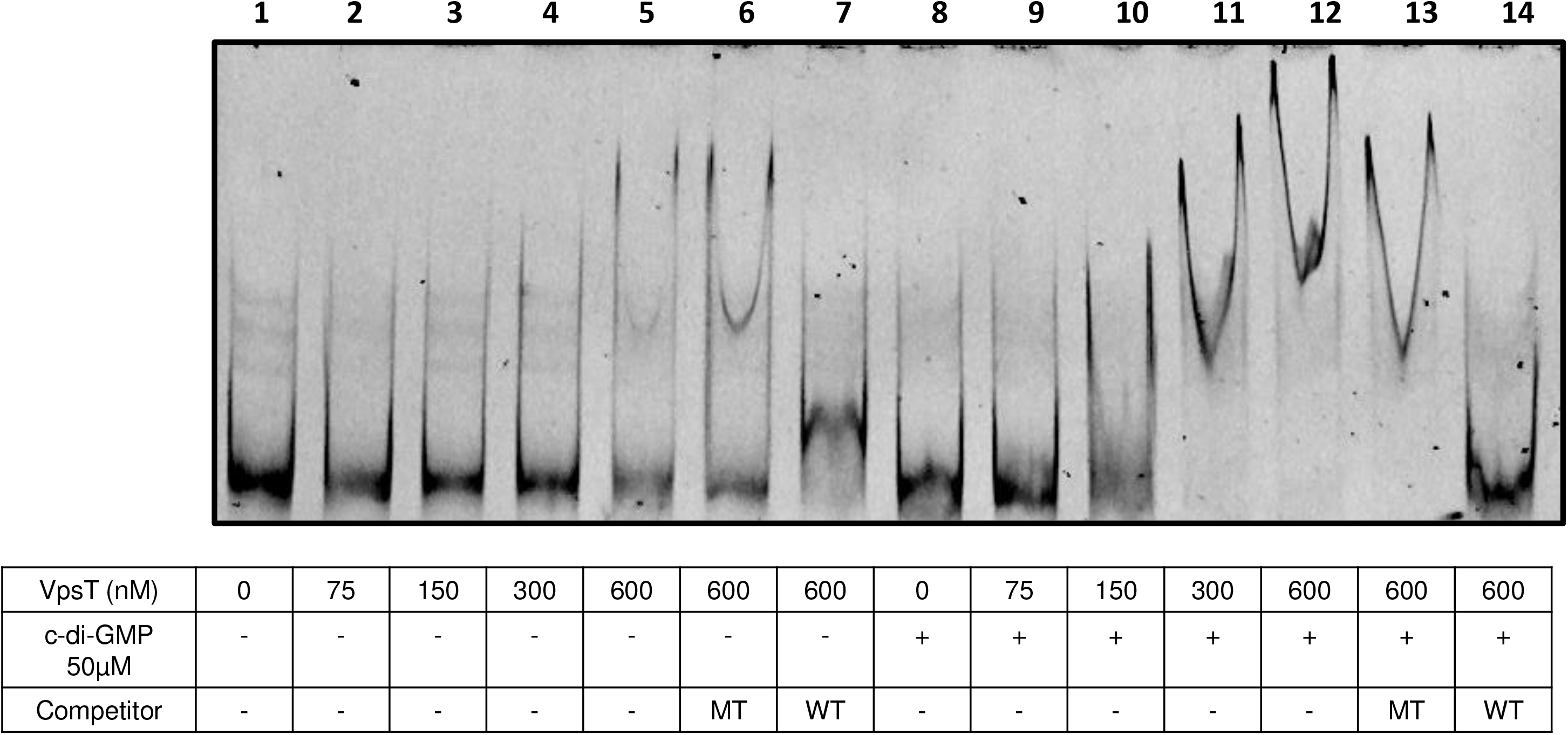
C-di-GMP enhances VpsT interaction with the *katB* promoter in vitro. Increasing concentrations of VpsT-HIS were incubated with a FAM-labeled probe corresponding to the *katB4* promoter (Lanes 1-5). Unlabeled mutant (MT) or wild-type (WT) VpsT binding site competitor was added at 100X-molar excess relative to the labeled probe in reactions with 600 nM VpsT-HIS (Lanes 6,7). Lanes 8-14 are the same reaction conditions as Lanes 1-7 except 50 μM c-di-GMP was added to the binding reactions, as indicated by the + sign.

### C-di-GMP mediated HOOH survival is dependent on catalase

Since c-di-GMP increased *katB* expression and catalase activity, we hypothesized that high intracellular c-di-GMP would provide a survival advantage during H_2_O_2_ stress. We tested this hypothesis by measuring survival after H_2_O_2_ treatment in *V. cholerae* backgrounds (Δ*vpsL*) unable to make mature biofilms to specifically test if the transcription regulation of *katB* by c-di-GMP was responsible for any observed protection as opposed to matrix production or formation of multicellular biofilms (8, 10, 33, 34). *V. cholerae* expressing QrgB^Mut^ or QrgB were challenged with H_2_O_2_ and the gross culture viability (OD_600_) was monitored for three hours. We found that in the parent strain, expression of QrgB, but not QrgB^Mut^, led to significant protection from H_2_O_2_ stress (Figure 7A). To test this if this protection is dependent on c-di-GMP mediated catalase activity, we measured H_2_O_2_ survival in the Δ*katB*, Δ*katG*, and Δ*katB*Δ*katG* mutants. Consistent with our catalase activity results, deletion of *katB* in either the Δ*katB* and Δ*katB*Δ*katG* mutants displayed no c-di-GMP mediated H_2_O_2_ survival (Figure 7E and G) while survival of the Δ*katG* mutant during H_2_O_2_ treatment resembled the parent strain (Figure 7F).

**Figure 7.**
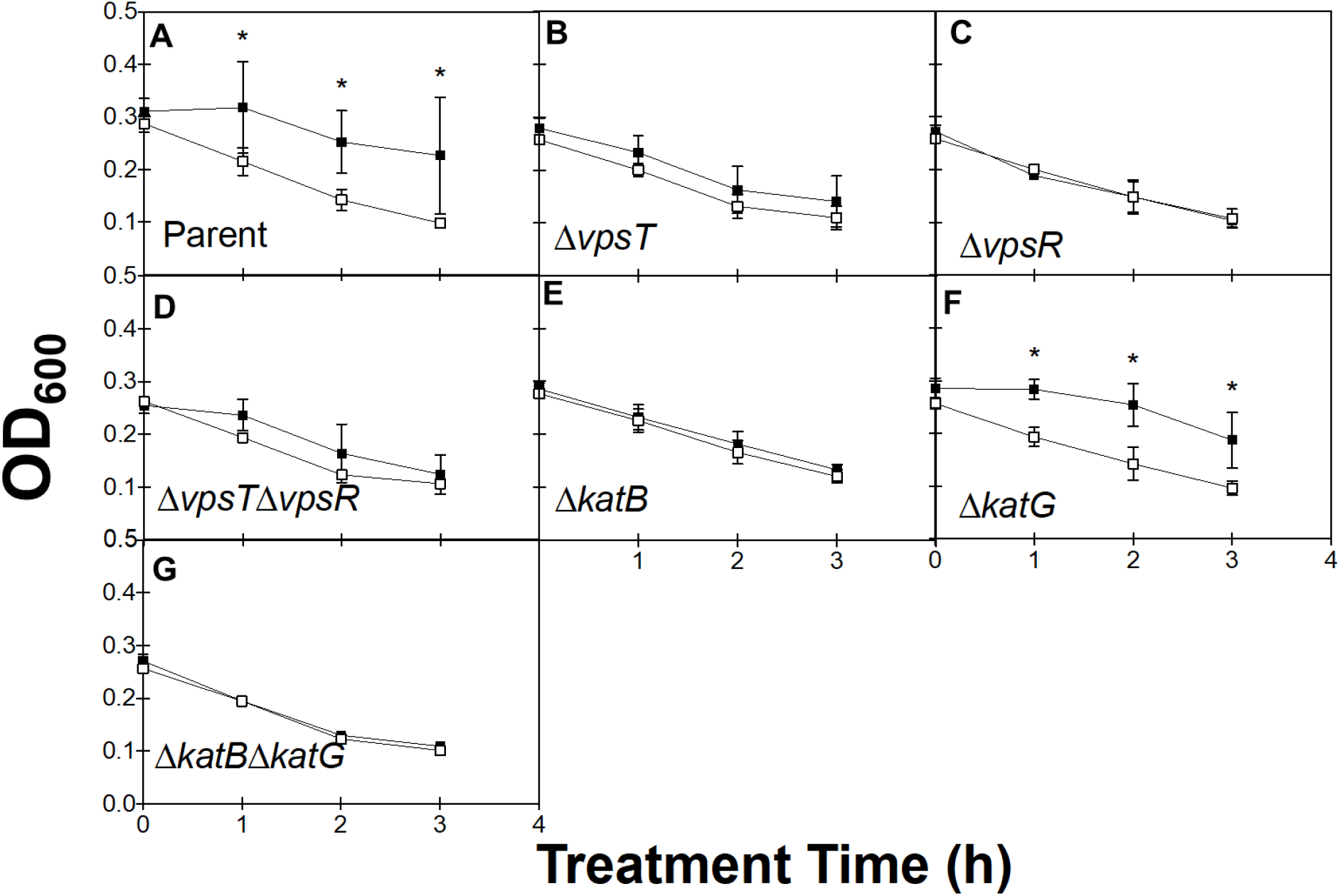
C-di-GMP enhances survival after H_2_O_2_ treatment. QrgB^Mut^ (open squares) or QrgB (squares) were induced with 100 μM IPTG in parent (A), Δ*vpsT*(B), Δ*vpsR* (C), Δ*vpsT*Δ*vpsR* (D), Δ*katB* (E), Δ*katG* (F), and Δ*katB*Δ*katG* (G)strains until an OD_600_ of 0.3. H_2_O_2_ was added to cultures at a final concentration of 12.5 mM. Cell death was monitored by measuring OD_600_ every hour after H_2_O_2_ addition (Time 0). * Indicate a p < .05 when compared to the untreated control at that timepoint determined by Two-Way ANOVA followed by Tukey’s multiple comparisons test. Bars are averages from three biological replicates and error bars indicate standard deviation.

As *vpsT* and *vpsR* were necessary for c-di-GMP dependent induction of *katB* expression and catalase activity (Figure 1 and 4), we hypothesized deletion mutations of these transcription factors would decouple c-di-GMP signaling from survival during H_2_O_2_ treatment. Indeed, strains lacking *vpsT*, *vpsR*, or both *vpsT* and *vpsR* lost c-di-GMP-mediated survival during H_2_O_2_ treatment (Figure 7B-D). Taken together, these data suggest c-di-GMP increases *katB* expression and KatB catalase activity through the c-di-GMP dependent transcription factors *vpsT* and *vpsR* resulting in increased survival during H_2_O_2_ treatment (Figure 8).

**Figure 8.**
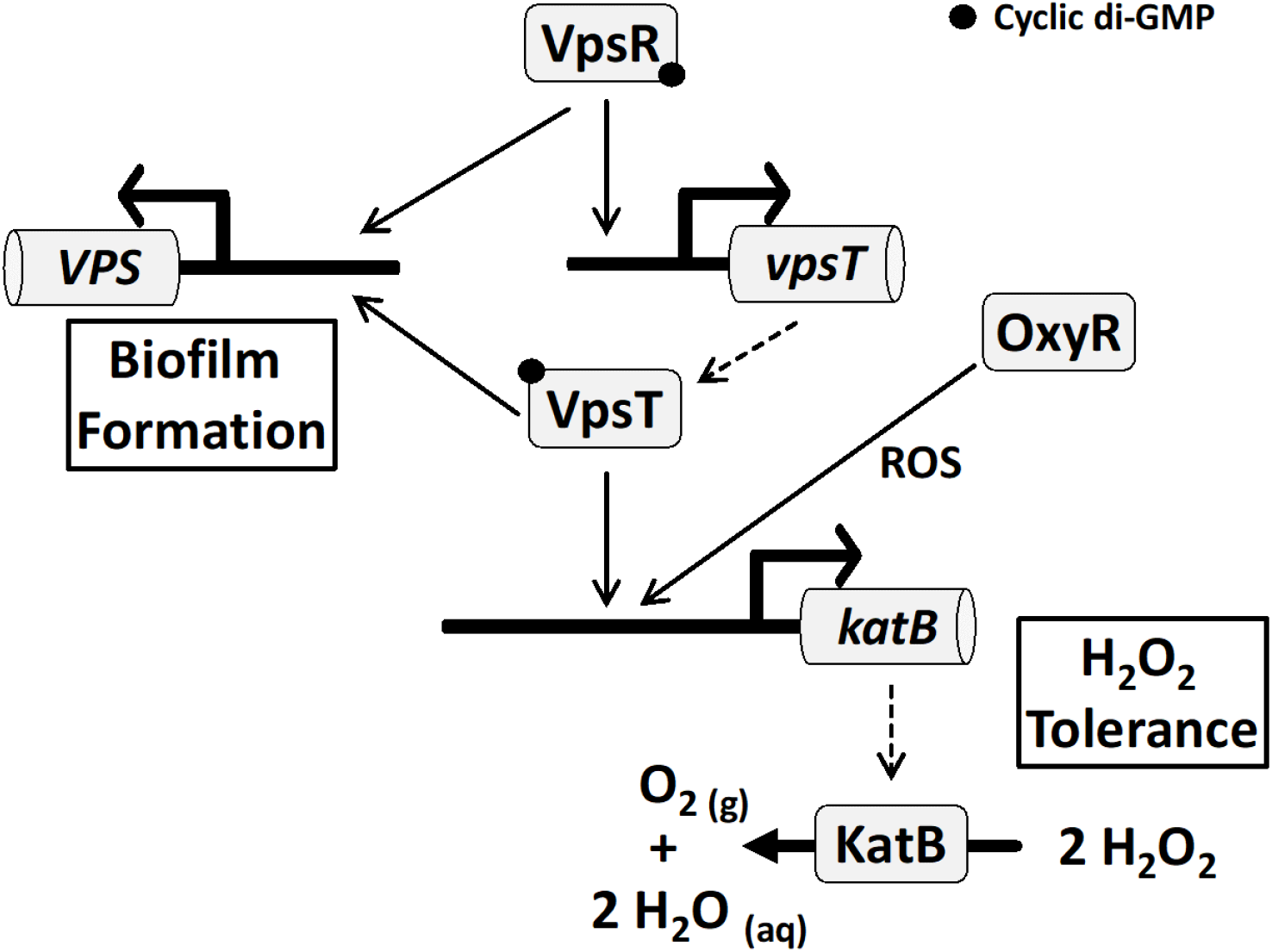
Model for the c-di-GMP regulatory network controlling biofilm formation and H_2_O_2_ tolerance. In response to high intracellular c-di-GMP, VpsR becomes activated and induces expression of *vpsT*. VpsT, in turn, is activated by c-di-GMP and both VpsR and VpsT activate genes in the *vps* operons to promote biofilm formation. Our data indicates c-di-GMP activated VpsT also induces expression of *katB* which increases catalase activity of the population which is able to turn over H_2_O_2_ and promote survival after H_2_O_2_ treatment. Our data also demonstrates induction of *katB* by H_2_O_2_ occurs independently of the c-di-GMP signaling network and is likely controlled by OxyR, a transcription factor involved in a broader antioxidant production program.

## Discussion

In this work, we sought to determine if c-di-GMP provides resistance to ROS in *V. cholerae*. Using a plasmid-based system to modulate intracellular c-di-GMP, we observed increased gas production after the addition of H_2_O_2_, suggesting c-di-GMP positively influenced catalase activity. We determined that *katB* was responsible for c-di-GMP regulated catalase activity and found that *katB* transcription was increased 5-fold in the parent, but this induction was lost in strains deficient for the c-di-GMP dependent transcription factors VpsT and VpsR. Measuring *katB* transcription in a heterologous host revealed that VpsT was sufficient to induce *katB* expression under high c-di-GMP conditions, and in vitro DNA binding assays demonstrated VpsT specifically bound to the *katB* promoter in a c-di-GMP dependent fashion. Lastly, we showed that c-di-GMP mediated survival after H_2_O_2_ treatment was dependent on *vpsT, vpsR*, and *katB*.

In agreement with our findings, other groups have observed an increase in *katB* expression under certain biofilm inducing conditions which modulate intracellular c-di-GMP, such as incubation with norspermidine (35). Additionally, transcriptomic data from experiments where c-di-GMP or VpsT was artificially induced in *V. cholerae* suggested *katB* expression was a positively regulated by c-di-GMP (13, 36). Interestingly, c-di-GMP and VpsT were shown to down-regulate production of the stationary phase sigma factor RpoS, in turn decreasing survival to various stressors including H_2_O_2_, which is in contrast to our results (Figure 6A) (20). These experiments were done with the El Tor biotype strain C7258, which is part of the serogroup Ogawa while the strain used in the current study (C6076 str2) is part of the serogroup Inaba. While there are no studies describing differences in transcriptional regulation between the two serogroups, we hypothesize there may be serogroup dependent differences in c-di-GMP signaling, which would explain the contrasting results.

Unlike the Δ*vpsT*Δ*vpsR* double mutant, c-di-GMP was able to induce the express of the *katB* reporter in the single *vpsT* and *vpsR* deletion background, albeit the difference was not statistically significant. However, only production of VpsT and c-di-GMP, but not VpsR and c-di-GMP, increased expression of the *katB* promoter in *E. coli* (Figure 4). C-di-GMP mediated catalase activity and survival in H_2_O_2_ were also lost in both the Δ*vpsT and* Δ*vpsR* backgrounds. Thus, our evidence suggests that phenotypes are primarily controlled by VpsT (Figure 8). VpsR is required for c-di-GMP mediated induction of *vpsT* in *V. cholerae*, and thus we conclude it has an indirect effect on *katB* transcription (31) (Figure 8).

A predicted VpsT binding site was previously reported approximately 150 bps upstream of the ATG start codon (37). Our results indicate that even when the VpsT binding site was present in the *katB4* transcriptional fusion, induction by c-di-GMP was lost. We note that the predicted VpsT sequence was found in the template strand while predicted and validated VpsT binding sites in the *vpsL* and *vpsA* promoters were found in the coding strand (32, 37). Whether this difference in strand binding alters VpsT transcriptional regulation warrants further investigation.

Although our work focuses on *V. cholerae*, the association between c-di-GMP and protection against ROS has been shown in other bacteria. In *Listeria monocytogenes*, deletion of genes encoding PDE domains have elevated biofilm production and H_2_O_2_ tolerance (9). However, it is not known whether H_2_O_2_ tolerance was caused by increased EPS production, antioxidant production, or both. In *Mycobacterium smegmatis*, a relative of the human pathogen *M. tuberculosis*, H_2_O_2_ stimulates production of intracellular c-di-GMP, which inactivates the transcriptional repressor HpoR. As a result, ROS defense genes are up-regulated and increase H_2_O_2_ tolerance (30). This differs from our work in two facets: first, H_2_O_2_ does not act as a first messenger that stimulates c-di-GMP activity in *V. cholerae* (Figure S2) and second, VpsT and c-di-GMP regulate a wide array of genes whereas HpoR-c-di-GMP has been shown to only regulate expression of the *hpoR* operon. Despite the differences in regulation, the connection between c-di-GMP and ROS tolerance is evident in bacteria from diverse phylogenetic backgrounds.

## Acknowledgments

This material was supported by NIH grants GM109259, GM110444, and AI130554 awarded to CMW. We would like to thank the College of Natural Science and the Department of Microbiology and Molecular Genetics at Michigan State University for the University Enrichment Fellowship and the Rudolph Hugh and Bertina Wentwoth Fellowship awarded to NLF. We would also like to thank Geoffrey B. Severin and Brian Y. Hsueh for critically reading and providing comments on the manuscript.

